# Gene expression profiling of Tea (*Camellia sinensis* L.) in response to biotic stress using microarrays

**DOI:** 10.1101/2020.03.17.994806

**Authors:** S. Ashokraj, E. Edwin Raj, K.N. Chandrashekara, R. Govindaraj, T. Femlin Blessia, B. Radhakrishnan

## Abstract

The blister blight (BB) and grey blight (GB) diseases are the major biotic stresses, which affecting the plant health, yield and quality of tea. The study aims to understand the gene response of tea plants against destructing foliar diseases in terms of differential gene expression and their pathways through microarray analysis aid by MapMan^®^ software. The results of expression profile analysis showed that 235 in BB and 258 for GB genes were differentially expressed (at P<0.05) which involving in gene regulatory function as biotic stress response. Similarly, 76 and 86 differentially expressed genes involving in cellular response during BB and GB diseases, respectively. However, 28 in BB and 9 in GB differentially expressed (P<0.01) genes were putatively involved in biotic stress response. The study also identified differentially expressed 75 transcription factors (TFs) belongs to 23 TFs superfamily act as either transcriptional activators or repressors. The study helps to understand the differential gene expression pattern and its cellular, molecular and biological mechanisms of tea plants of two different diseases based on microarray analysis. Further studies using biotechnological tools on the stress-responsive genes in the germplasm may enable us for development of disease resistance.

## INTRODUCTION

Tea (*Camellia sinensis*) is a common non-alcoholic beverage consumed worldwide in across all groups of people due its aroma, taste and medicinal properties. Tea manufacturing is depends young leaves due to high accumulation of secondary metabolites such as polyphenols, catechins and polysaccharides. Growth and yield of tea plant affected by various biotic/abiotic stresses besides changing climate. Blister blight (BB) and grey blight (GB) are the major foliar diseases leads to crop loss up to 40% and 20%, respectively, in tea plantation and decreasing made tea quality by affecting polyphenol and catechins contents (Jayaswall et al., 2016; Joshi et al., 2009). BB is caused by *Exobasidium vexans* Massee (Gadd and Loos, 1948), is an obligate biotrophic plant pathogen belongs to basidiomycete and GB is caused by *Pestalotiopsis theae* (Sanjay et al., 2008). Currently, management of the diseases are highly depending on synthetic chemical fungicides. However, conventional chemical practices are creating major issues which including phytotoxicity, fungicide resistance and accumulation of residues in made tea. Moreover, alternative fungicides biocontrol agents are developed which also having certain limitation for application at field level. Likewise, development of disease resistant varieties through existing methods is time consuming process due to heterozygous and self-incompatibility nature of tea besides identification resistant accessions from the population (Jayaswall et al., 2016). However, high economic threshold levels by the biotic stress insisting the researches on plant-pathogen interaction studies and identification of stress-responsive genes for the development of disease resistant plants.

Therefore, incorporation of modern molecular and biotechnological approaches along with conventional breeding approaches would be given effective strategy in development of disease resistant tea clones. The functional genomics and gene expression analysis would elucidate the stress-responsive genes, which plays major role in marker assisted selection (MAS) breeding. Functional genomics is used to elucidate biological activities of large set of genome information through gene expression analysis. Microarray technology is being effectively utilised for comprehensive and simultaneous gene expression profiling of the plant (Lodha and Basak, 2012). Incorporation of these gene expression profiling with pathway and bioinformatics analysis will explain the molecular and cellular mechanisms of host-pathogen interaction and defense signalling pathways during the infection period (Wan et al., 2002). Microarray technology is successfully employed for studies on large scale of genes and their expression at a time. The quantitative results of gene expression resulting the qualitative changes to regulatory processes from cellular to organism level (Xiang et al., 2003). To fill the gap of plant responses to biotic stress of two different diseases in tea, the study aims to elucidate the expression profile of biotic stress-responsive genes and transcription factors (TFs).

## MATERIALS AND METHODS

### Plant materials and biotic stress conditions

For the microarray analysis, biotic stress treatment was imposed in two-year-old plants at the UPASI TRF Experimental Farm, Valparai. Fungal spores were collected from BB and GB infected plants and fungal spores of respective organisms were confirmed under microscope. The spore suspensions were prepared for both *E. vexans* (10^6^) and *P. theae* (10^5^) in sterile distilled water and the inoculation was performed during 7:00 to 8:00 AM. The spore suspension was dropped on to surfaces sterilised, pre-wounded leaves i.e., 1^st^, 2^nd^ and 3^rd^ leaves of the plants. Control plants were inoculated with sterile water. The spores of *E. vexans* and *P. theae* were respectively inoculated in SA-6 and UPASI-10 cultivars which were maintained at 100% RH for 72 h in a shade and then transferred to glasshouse. Control and treated leaves were collected and stored in RNA Later (Invitrogen^®^) solution followed by the microarray analysis was performed at M/s. Genotypic Technology Pvt. Ltd., Bangalore.

### Gene Expression profiling Using Agilent Platform

A 4 x 44K (AMADID: 043117) gene expression chip was designed with the probes having 60-mer oligonucleotides from mRNA sequences were downloaded from NCBI of *C. sinensis* and disease resistance genes of *Arabidopsis thaliana*. All the oligonucleotides were designed and synthesised *in situ* as per the standard methodologies of Agilent Technologies. Probes covering resistance genes were collected from Agilent catalog (AMADID 037661) and designed in Agilent eArray platform. BLAST was performed against the mRNA sequence databases to check the specificity of the probes.

### RNA Quality Control

RNA extraction was done using RNAqueous kit from Ambion according to manufacturer’s protocol. Total RNA integrity was assessed using RNA 6000 Nano Lab Chip on the 2100 Bioanalyzer (Agilent, Palo Alto, CA) and RNA purity was assessed by the NanoDrop^®^ ND-1000 UV-Vis Spectrophotometer (Nanodrop technologies, Rockland, USA).

### Labelling and microarray hybridization

The samples for gene expression were labelled using Agilent Quick-Amp labelling Kit (Part Number 5190-0442). A 500 ng of RNA samples were incubated with reverse transcription mix at 40 °C for cDNA synthesis primed by oligo-dT with a T7 polymerase promoter. The cleaned up double stranded cDNA were used as template for cRNA generation. cRNA was generated by *in vitro* transcription and the dye Cy3 CTP (Agilent) was incorporated during this step. The cDNA synthesis and *in vitro* transcription steps were carried out at 40°C. Labelled cRNA was cleaned up and quality assessed for yields and specific activity.

### Hybridisation and scanning

The labelled cRNA samples were hybridised on to an Agilent Platform custom designed *C. sinensis* 4×44K and 1650 ng of cy3 labelled samples were fragmented and hybridised. Fragmentation of labelled cRNA and hybridisation were done using the Gene Expression Hybridisation kit of Agilent (Part Number 5188-5242). Hybridisation was carried out in Agilent’s Surehyb Chambers at 65 °C for 16 h. The hybridised slides were washed using Agilent Gene Expression wash buffers (Part No: 5188-5327) and scanned using the Agilent Microarray Scanner G2505C at 5 micron resolution.

### Microarray data analysis

Data extraction from images was done using Feature Extraction software v 10.7 and analysed using GeneSpring GX version 11.5 software (Agilent). Normalization of the data was done in GeneSpring GX using the 75^th^ percentile shift and normalization to specific samples. Differentially regulated genes were clustered using hierarchical clustering to identify significant gene expression patterns.

### MapMan Ontology mapping

In order to identify functionally related genes and get pictorial representations for the cellular, molecular and biotic stress response pathways BIN structures of MapMan^®^ representation ontology was adopted (Rotter et al., 2007). The original BIN assignments for *Arabidopsis thaliana* were based on publicly available gene annotations from TIGR (The Institute for Genomic Research) using a process which involved alternation between automatic recruitment and manual correction.

## RESULTS AND DISCUSSIONS

The symptom of disease was noticed after 3-4 day of inoculation as spot grow in size. The morphological and microscopic observation were confirmed the BB and GB diseases on the respective cultivars. The BB disease initially developed an oil spot to circular stage after 11 days of inoculation. Likewise, GB disease development also observed after 15 days.

### Identification of differentially expressed genes to biotic stresses

The 4×44K array comprised of 45220 features including 43803 probes and 1417 Agilent controls. *Camellia* genes: For 758 fungal resistance genes, on an average of one probe per gene were designed in Agilent eArray platform. *Arabidopsis* genes: Probes covering 27336 genes were collected from Agilent catalogue (AMADID 037661). In addition, For 230 disease resistance genes probes were designed. Finally, 28338 probes were designed and 15465 specific probes were replicated to fill the remaining spots. The data were archived at the public microarray database under NCBI accession number GSE45360.

The differential expressed genes were determined by their expression of more than twofold (up regulation) or less than halffold (down regulation) (P<0.05) during biotic stress conditions. In both the diseases, more number of genes were differently expressed during infected condition, which showed that the plants have dynamically responded during biotic stresses. In BB, of the 1382 differently expressed genes 789, 216 and 377 were expressed during infection, control and both the conditions, respectively. In case of GB, of the 2077 differently expressed genes 1478, 168 and 431 were expressed during infection, control and both the conditions (Fig. 1). Further, these genes were used for analysis of gene regulation, cellular and stress response during disease infection.

**Fig. 1:**
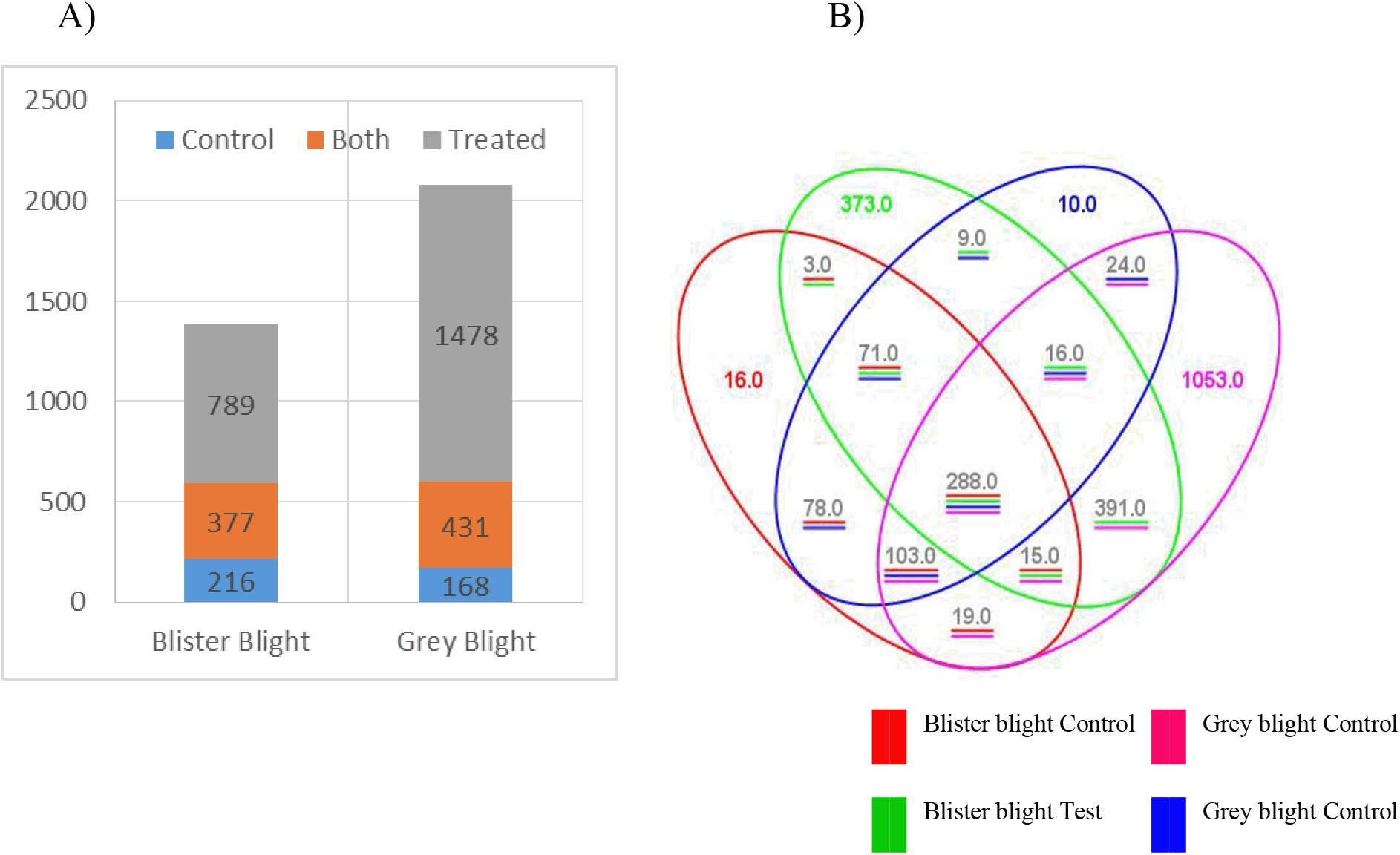
Illustration of overall differential expression during BB and GB disease transition (A); and Distribution of differentially expressed genes in control and infected plants (B).

### Gene regulation during biotic stress conditions

The expression patterns of gene regulation was depicted for both BB and GB disease using MapMan^®^ (Fig. 2). This analysis MapMan bioinformatic tool indicated a biotic stress induced enrichment in genes related to biotic and abiotic stress responses, including proteinase inhibitors, putative stress-induced proteins, disease resistance-related proteins, pathogenesis-related proteins, defense signalling and heat shock proteins. Overall, 127 and 108 genes were up and downregulated, respectively during BB infection. Likewise, 117 and 141 genes were up and downregulated, respectively during GB infection (Table 1). The results indicated that the stress-responsive genes are regulating defense signalling pathway in the plants to the respective biotic stresses (Jayaswall et al., 2016; Li et al., 2017).

**Fig. 2:**
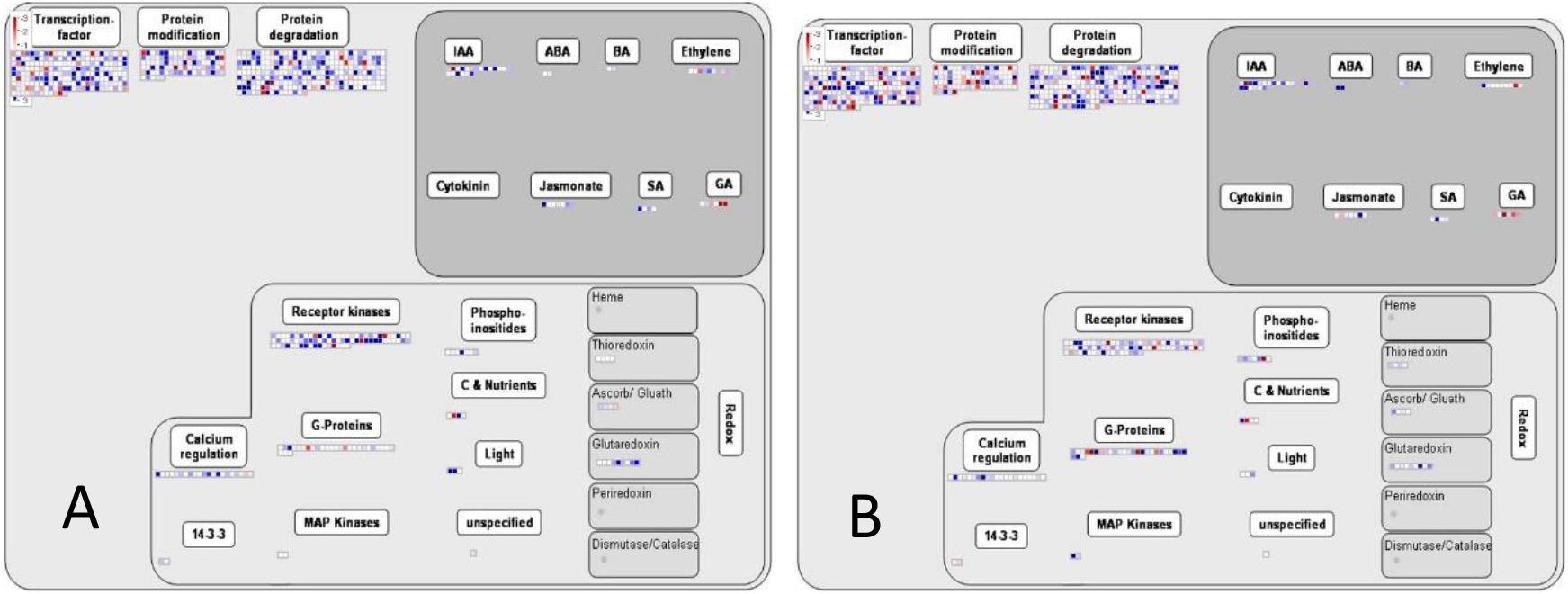
An overview of gene regulation during biotic stress (A) BB and (B) GB.

**Table-1.** Differential expression of gene regulation during biotic stress condition

| **Gene regulatory function** | **BBUR** | **BBDR** | **GBUR** | **GBDR** |
| --- | --- | --- | --- | --- |
| Transcription factors | 32 | 38 | 32 | 42 |
| Protein modification | 17 | 16 | 9 | 27 |
| Protein degradation | 44 | 28 | 44 | 45 |
| IAA | 5 | 2 | 6 | 2 |
| ABA | 0 | 0 | 2 | 0 |
| Ethylene | 0 | 2 | 1 | 1 |
| Jasmonate | 1 | 0 | 1 | 0 |
| SA | 1 | 0 | 1 | 0 |
| GA | 0 | 3 | 0 | 4 |
| Receptor kinase | 17 | 10 | 10 | 12 |
| Phosphoinositides | 1 | 0 | 0 | 1 |
| C & Nutrients | 1 | 1 | 1 | 1 |
| Calcium regulation | 3 | 2 | 2 | 0 |
| G-Protein | 1 | 5 | 6 | 6 |
| MAP Kinases | 0 | 0 | 1 | 0 |
| Light | 2 | 0 | 0 | 0 |
| Ascorb/Gluath | 0 | 1 | 0 | 0 |
| Glutaredoxin | 2 | 0 | 1 | 0 |

BBUR – upregulated genes during BB infection; BBDR - downregulated genes during BB infection; GBUR - upregulated genes during GB infection; GBDR - downregulated genes during GB infection.

### Cellular responses in tea leaves during biotic stress conditions

MapMan^®^ overview of cellular response during both diseases were depicted in Fig. 3. Overall, 36 and 40 cellular responsive genes were up and downregulated, respectively during BB infection. Likewise, 39 and 47 genes were up and downregulated, respectively during GB infection (Table 2). The differentially expressed genes of cellular functions indicated that the tea plant produced cellular or organismal gene products that could be responsible for plant-pathogen interactions and defense signalling (Wan et al., 2002). Further molecular studies will explain complete cellular functions during biotic stress.

**Fig. 3:**
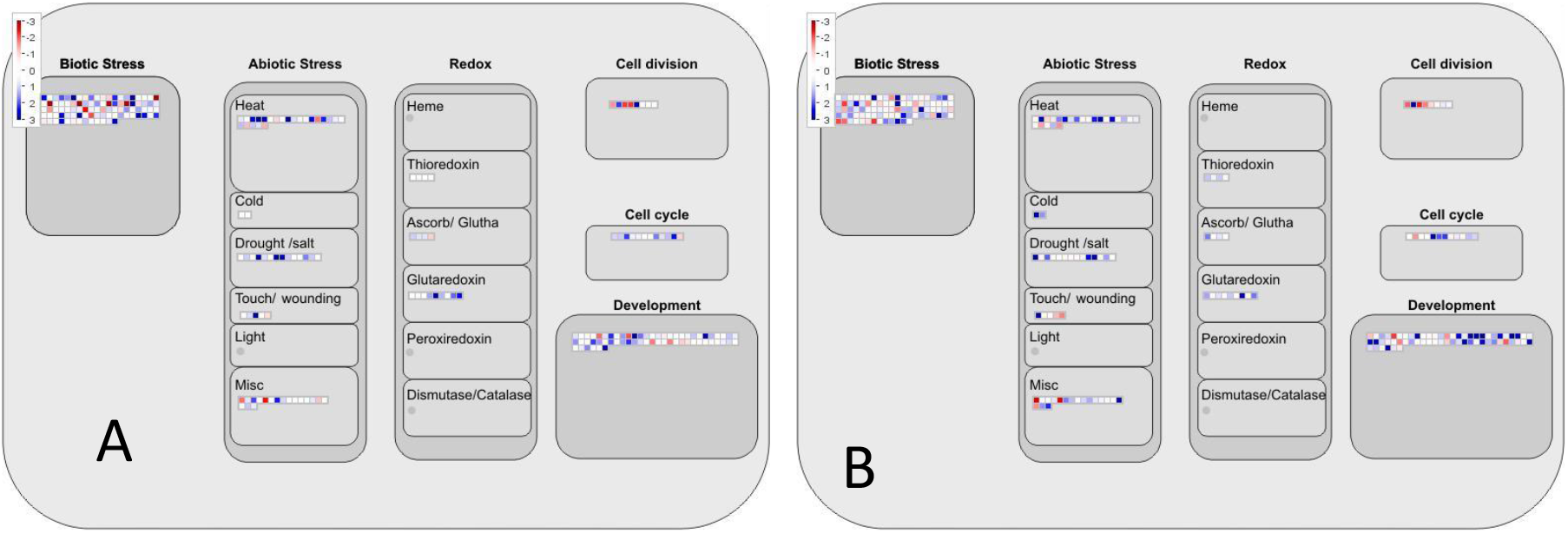
Overview of cellular responses in tea to biotic stress (A) BB and (B) GB.

**Table 2.** Differential expression of cellular response genes during biotic stress condition

| **Cellular response** | **BBUR** | **BBDR** | **GBUR** | **GBDR** |
| --- | --- | --- | --- | --- |
| Biotic stress | 14 | 19 | 9 | 21 |
| Heat | 6 | 4 | 5 | 4 |
| cold | 0 | 0 | 1 | 0 |
| Drought/salt | 3 | 0 | 3 | 0 |
| Touch/wound | 1 | 0 | 1 | 2 |
| Unspecified | 2 | 3 | 2 | 3 |
| Ascorbate/Gluthathione | 0 | 1 | 0 | 0 |
| Glutaredoxin | 0 | 2 | 1 | 0 |
| Cell Division | 2 | 3 | 1 | 4 |
| Cell cycle | 2 | 1 | 2 | 2 |
| Development | 6 | 7 | 14 | 11 |

Footnotes are same as table-1

### Differential expression profiles during biotic stress conditions

Over all pathways and their significant gene expression were assumed using MapMan (Fig. 4). For identification of highly significant differential expressed genes p < 0.01 used as a cut-off value. Overall, 28 genes were downregulated during BB infection. Likewise, 4 and 5 genes were up and downregulated, respectively during GB infection (Table 3). The higher number of downregulated genes during BB infection indicated that these genes are plays a role in plant-pathogen interaction. Further molecular studies will explore their role in BB infection. Overall pathway depicting the defense signalling and response of the plants during the biotic stress conditions. The molecular approaches on these genes will be useful for pathway engineering for development of resistant plants (Kries and O’Connor, 2016).

**Fig. 4.**
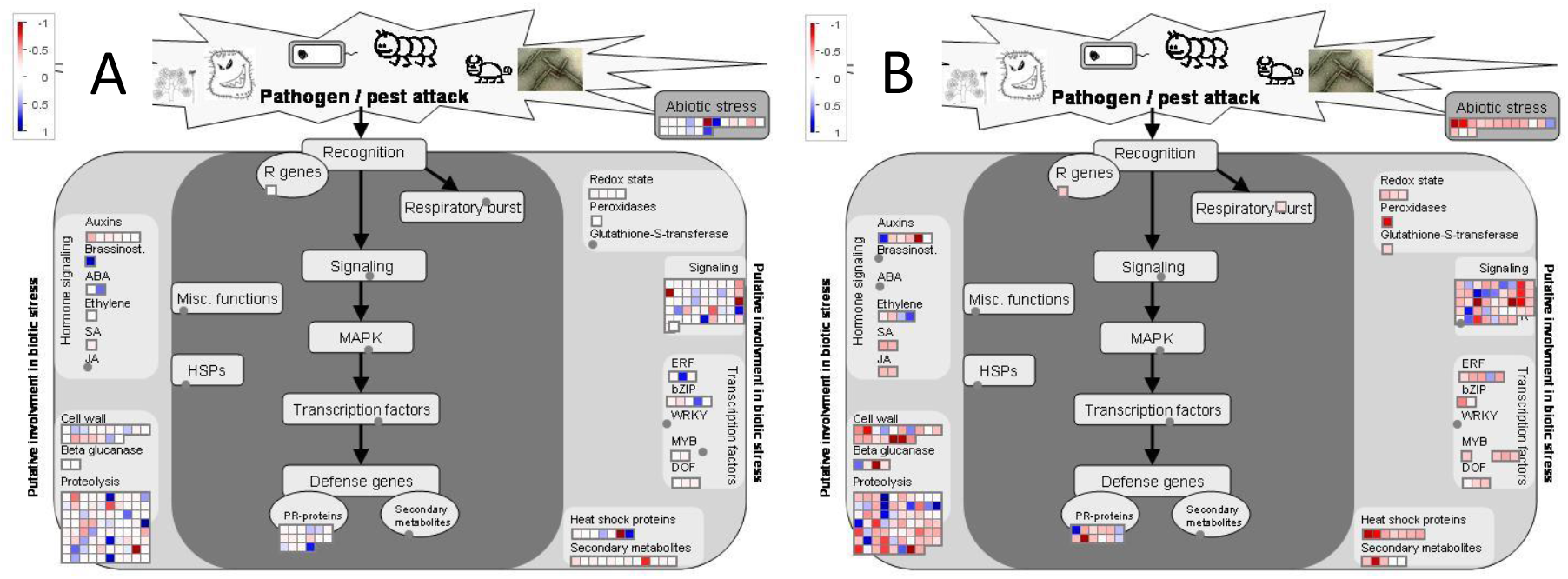
Genes that were shown to be differentially expressed using p < 0.01 as a cut-off value (visualised by MapMan). A) BB and B) GB.

**Table 3.**
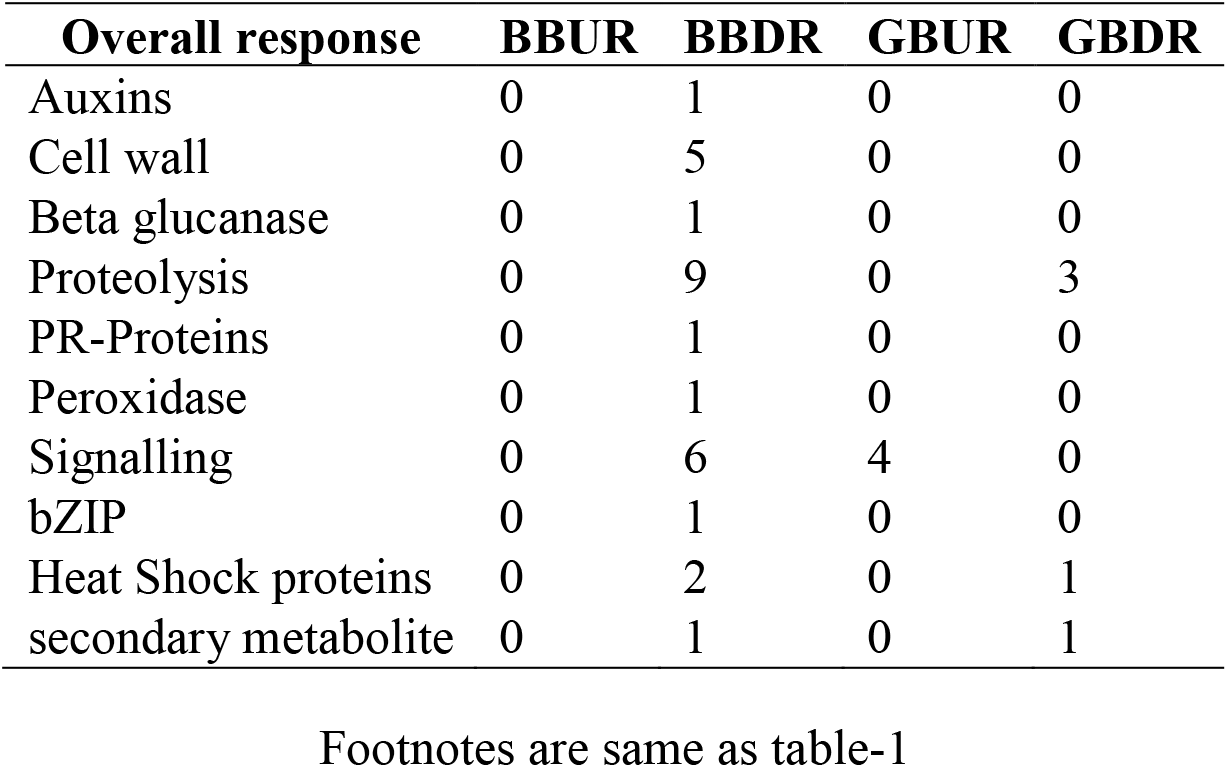
Differentially expressed disease responsive genes using p < 0.01 as a cut-off value

### Expression profiling of Transcription factors (TFs)

Microarray analysis showed that there were 75 TFs differently expressed during biotic stress. Out of 40, 19 and 21 were down and upregulated, respectively during BB infection. Out of 35 TFs, 16 and 19 were down and upregulated, respectively during GB infection (Table 4). Twenty-three TF superfamilies were screened, in which 11-12 TFs were downregulated and 13-14 TFs were upregulated. Amongst the 66 TFs, Dof, ERF, NAC and WRKY were important TFs play a vital role in the control of hormone signalling and pathways, secondary metabolic processes, cell cycle regulation, defense and wound repair mechanisms, activation of PR genes and multiple defense responses (Li et al., 2015). Up and downregulation of Dof TFs during GB showing that there was a alterations in the secondary metabolic processes, such as the biosynthesis of glucosinolates and flavonoids, phytochrome and cryptochrome signalling, and cell cycle regulation. Due to up-and-downregulation of many ERF TFs in both BB and GB showing that, the plants were unable to coordinate stress signalling with the activation of defense mechanisms, which leads to the development of the disease. Upregulation of NAC TFs indicated that the proteolytic activation of a plasma membrane binding through the promotors of PR genes by pathogens. Downregulation of WRKY superfamily during BB infection revealed that during pathogenesis the multiple defense responses of the plant was repressed especially in sensing pathogen-associated molecular patterns (PAMP) in downstream of mitogen-activated protein kinase (MAPK) cascades (Jayaswall et al., 2016; Li et al., 2015)

**Table 4.**
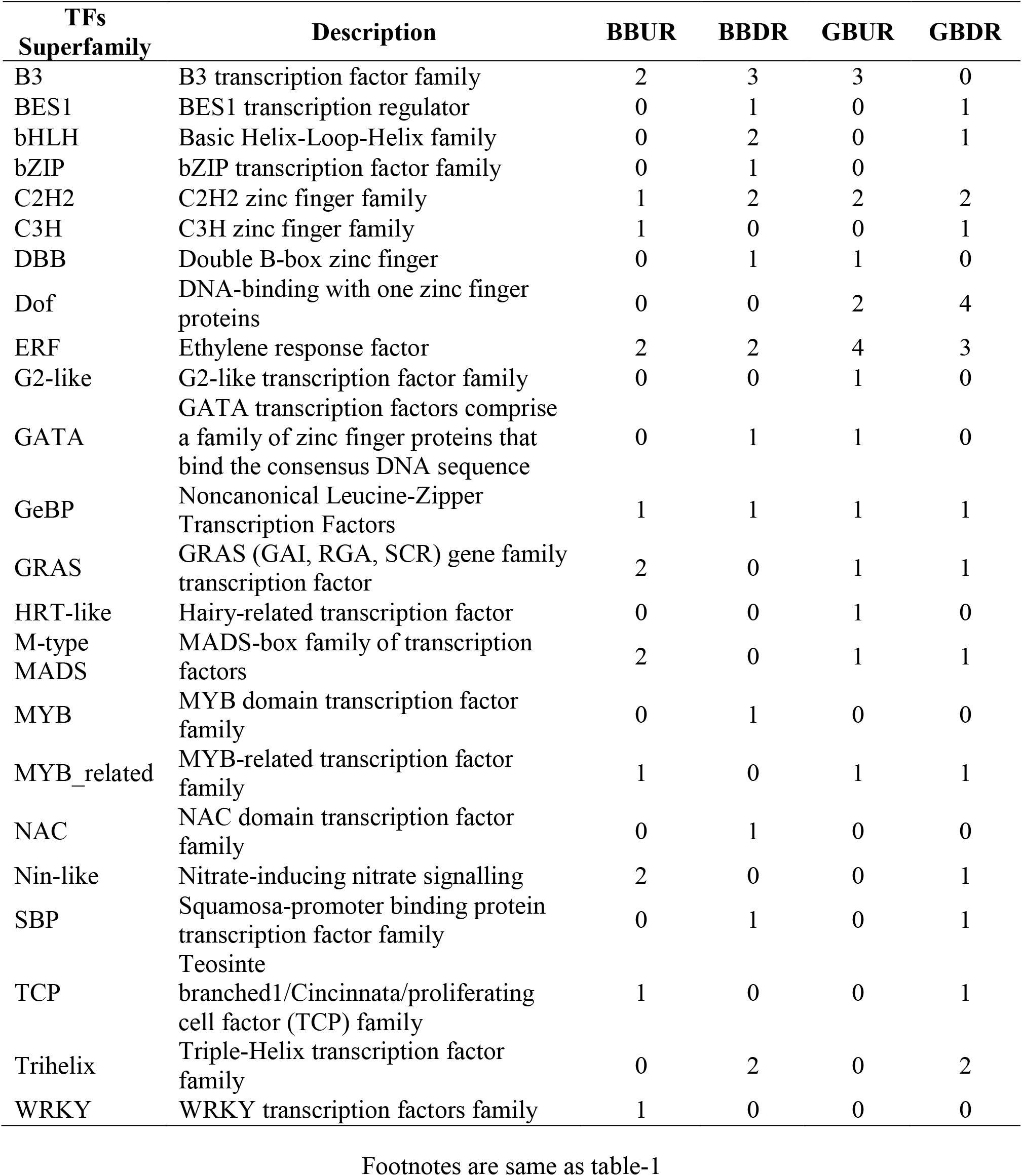
Differential expressed TFs during BB and GB infection.

## CONCLUSION

The study found MapMan implementation for tea beneficial as it would facilitate biological interpretation, support an ontology-based statistical data analysis and provide users a global overview of the results. From microarray analysis, we have identified certain biotic stress responsive genes related to gene regulation, cellular function and signalling pathways. Further molecular and biotechnological studies will provide the way to develop the resistance plants to BB and GB disease. The stress-responsive genes also utilized for MAS breeding.

## Acknowledgement

The authors are thankful to Tea Board of India for the financial support to conduct this study. The authors are also grateful to UPASI Tea Research Foundation, Tea Research Institute, Valparai for providing facilities for this study.

